# Training data composition affects performance of protein structure analysis algorithms

**DOI:** 10.1101/2021.09.30.462647

**Authors:** Alexander Derry, Kristy A. Carpenter, Russ B. Altman

## Abstract

The three-dimensional structures of proteins are crucial for understanding their molecular mechanisms and interactions. Machine learning algorithms that are able to learn accurate representations of protein structures are therefore poised to play a key role in protein engineering and drug development. The accuracy of such models in deployment is directly influenced by training data quality. The use of different experimental methods for protein structure determination may introduce bias into the training data. In this work, we evaluate the magnitude of this effect across three distinct tasks: estimation of model accuracy, protein sequence design, and catalytic residue prediction. Most protein structures are derived from X-ray crystallography, nuclear magnetic resonance (NMR), or cryo-electron microscopy (cryo-EM); we trained each model on datasets consisting of either all three structure types or of only X-ray data. We find that across these tasks, models consistently perform worse on test sets derived from NMR and cryo-EM than they do on test sets of structures derived from X-ray crystallography, but that the difference can be mitigated when NMR and cryo-EM structures are included in the training set. Importantly, we show that including all three types of structures in the training set does not degrade test performance on X-ray structures, and in some cases even increases it. Finally, we examine the relationship between model performance and the biophysical properties of each method, and recommend that the biochemistry of the task of interest should be considered when composing training sets.

## 1. Introduction

Understanding the complex interaction between protein structure and function is crucial for elucidating disease mechanisms, discovering new drug treatments, and many other key questions in biology and medicine. The advent of machine learning techniques for analysis of 3D protein structures has led to great advancements in many tasks across structural biology, including protein structure prediction,^1–3^ protein-protein and protein-ligand interface prediction,^4,5^ and the design of novel protein sequences and structures.^6–8^

Despite the strong performance of these machine learning methods, the degree to which this performance will be replicated on new data depends on a match between the distribution of structures used for training and the distribution of structures to which the methods are applied. In the case of structural biology, the distribution of the underlying data (the actual 3D positions of atoms) is governed by the laws of physics and is well-characterized. However, the data provided to computational algorithms are not direct measurements of these positions. Instead, input data consists of 3D structures derived from X-ray crystallography, NMR and cryo-EM experiments. The fundamental differences between these technologies result in 3D atomic structures with measurably different underlying distributions.^9,10^

Multiple studies^11,12^ have characterized these differences. X-ray crystallography generally produces structures with higher atomic resolution than NMR^13^ and is better at solving covalent geometry and torsion angles. However, it can only produce structures for crystallizable molecules, and does not fully capture dynamics in solution. This experimental limitation results not only in different structures for a single protein, but also different distributions of protein families within the sets of structures solved by each method. Conversely, NMR structures reflect dynamics and protein structure in solution, but are based on ensembles of noisier local measurements and have lower resolution. NMR also tends to be used on smaller proteins. Cryo-EM has been increasingly used for large proteins and has several logistical advantages.^14^ However, resolution is generally low and its application can be hampered by technical considerations. Cryo-EM measures electron potential rather than electron density, a distinction that leads to different structural details.^15^ Thus, the three primary structural determination methods have a complex set of possible biases which are not well characterized in the context of machine learning. However, by empirically examining the structures generated and the algorithms that use them for training, we can better understand the magnitude and importance of these differences.

The increasing prominence of cryo-EM (Figure 1) makes this issue more pressing, as it is shifting the distribution of data more rapidly, causing the potential for model bias due to dataset shift.^16^ X-ray crystallography has been the dominant method of structure determination for proteins deposited in the Protein Data Bank (PDB) for decades,^17^ so most analytic methods developed have taken one of two approaches: removing all non–X-ray structures, resulting in a loss of increasingly valuable NMR and cryo-EM data; or including all data regardless of experimental method, which assumes that there is not a meaningful difference across methods. We hypothesize that the experimental source of protein structure data is important to consider when constructing training datasets and evaluating machine learning algorithms, especially as such models are increasingly being used to guide biological experimentation.

**Fig. 1.**
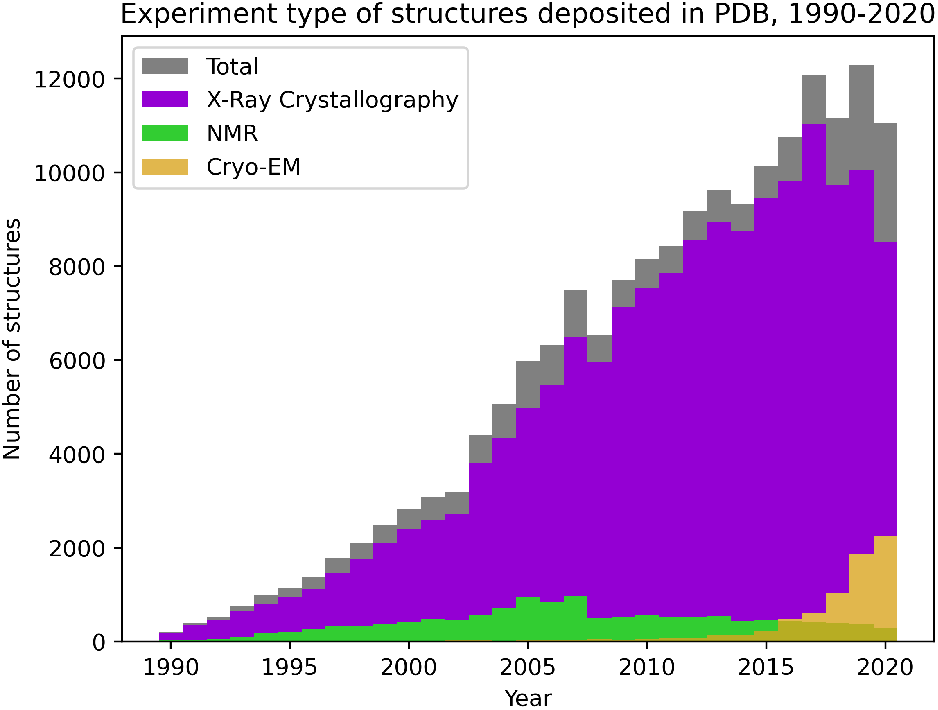
Change in distribution of structures deposited to the PDB from 1990-2020.

In this study, we explore the effect of training set composition on the performance of different algorithms applied to 3D protein structures. We focus on three common tasks: estimation of model accuracy (EMA), protein sequence design, and catalytic residue prediction. For each task, we use publicly available implementations of machine learning methods which learn directly from 3D structure. Each method is trained on datasets containing either (1) X-ray data only or (2) a combination of X-ray, NMR, and cryo-EM data. We evaluate the performance of each of these models trained on different data using a fixed test set containing all three types of structures. Finally, we provide preliminary recommendations to maximize the performance of structure-based algorithms in a deployment setting.

Code is available at https://github.com/awfderry/ml-structure-bias and datasets can be downloaded at https://zenodo.org/record/5542201.

## 2. Methods

### 2.1. Experimental Design

We choose three tasks spanning a variety of applications and describe each here, along with the evaluation metrics used to assess performance.

**Estimation of model accuracy** is the task of scoring how close a proposed protein structure (called a “decoy”) is to the ground truth “target” structure. This is an important task for the blind evaluation of new structure prediction algorithms, and is thus a standard task in the biennial Critical Assessment of Structure Prediction (CASP) competition.^18^ The outputs are global and depend directly on the atomic structure of the target protein, so any model must learn the intricacies of this structure along with its potential biases.

#### Evaluation

Model accuracy is typically quantified using the Global Distance Test Total Score (GDT-TS), which is computed through the Local-Global Alignment (LGA) algorithm.^19^ Our models are trained to predict this quantity, and we evaluate performance using the Spearman rank correlation between predicted and actual GDT-TS for each decoy. We choose to use Spearman correlation for two reasons: (1) a major use case of model assessment algorithms is in the ranking of candidate structures for a specific target, and (2) the metric is insensitive to the specific values of GDT-TS, which can vary across targets. In addition to global correlation across all CASP targets, we report a mean per-target correlation which assesses the ability of a model to rank candidate models for an individual protein. Statistical significance between correlations is assessed using Fisher-transformed *z*-scores on the Pearson correlation coefficients (see Supplementary Data for details).

**Protein sequence design** is the inverse of the well-known structure prediction task— given a desired structure, the model produces an amino acid sequence predicted to fold into that structure. A successful protein design model will enable rapid generation of *de novo* structures that may, for example, tightly bind a desired ligand of interest or self-assemble into a nanomaterial without first requiring a natural protein that already demonstrates the activity of interest. This task requires a model to accurately learn sequence-structure relationships, and has recently been a popular task for deep learning models in structural biology.^20–23^

#### Evaluation

For evaluation, we follow the example of recent works^22,23^ and evaluate this task using perplexity and native sequence recovery. Perplexity is a common metric in natural language processing which represents the likelihood of a particular sequence occurring under the trained model, and recovery is the fraction of amino acids in the true sequence that are predicted by the model. Although these are proxy metrics, under the assumption that native sequences are optimized for their structures,^24^ we want to select for models that are better at assigning high probability to native sequences. Statistical significance between perplexity distributions on each dataset is assessed using Mann-Whitney U test (see Supplement).

**Catalytic residue prediction** involves predicting which amino acids in an enzyme are involved in functional activity, an important task for automatic annotation and active site identification. Many methods and databases use sequence-based models to predict function,^25,26^ but since function is closely tied to structure, it is important to approach functional residue prediction through a 3D lens. For this task, models must effectively learn structure-function relationships in situations that often rely on precise positions of critical atoms.

#### Evaluation

This task requires predicting binary labels for catalytic residues. Since this task is highly imbalanced (with far fewer functional residues than nonfunctional ones), we evaluate performance using the area under the precision-recall curve (AUPRC). Statistical significance is assessed using Mann-Whitney U test (see Supplement).

##### Algorithm selection

We trained one contemporary machine learning algorithm for each task in our experiments. We selected algorithms using the following criteria: (1) The model must be published and evaluated on the task in question. (2) The model must learn directly from 3D atomic coordinates, without the use of hand-crafted or derived structural features. (3) The model must report performance that is near state-of-the-art for the task, with some leniency due to variations in evaluation criteria. (4) Finally, the algorithm must be publicly available, including the ability to retrain on arbitrary protein structures from the PDB. This final point is crucial because our experiments require training a new model on each data split, while many published methods only include code for evaluating trained models on new data.

For both EMA and protein design, we use the recently published Geometric Vector Perceptron (GVP)^23^ graph neural network architecture, a method for learning from protein structure which has demonstrated state-of-the-art performance on both tasks as well as a variety of other problems in structural biology.^27^

For catalytic residue prediction we use DeepFRI,^28^ which uses a graph convolutional network to predict function protein function. The gradients of a DeepFRI model trained on enzyme classification can be used to identify catalytic residues with high accuracy. By adopting this method, we can evaluate not only predictions at residue level for catalytic site prediction, but also global predictions of enzyme class designated by the Enzyme Commission^29^ (EC).

##### Splitting methodology

For each task, we construct two separate training sets based on the two commonly used methodologies in the literature: one containing only structures solved by X-ray diffraction, and the other containing a mix of structures solved by X-ray, NMR, and cryo-EM. Validation sets were selected randomly from within each training set and were used to choose the final model. All evaluations for each task are conducted on a single held-out test set containing X-ray, NMR, and cryo-EM structures, and performance is evaluated separately for structures solved by each method. By using this setup, we mimic the scenario where published methods trained on varying input datasets are tested on new data which may be drawn from a different distribution. We also emphasize that we did not attempt to balance the distributions of experimental method in the training sets, or create an NMR-only or cryo-EM–only training set, since this would require either drastically reducing the training set size or oversampling NMR and cryo-EM structures, which would further bias our analysis.

### 2.2. Task-specific Methods

#### Estimation of Model Accuracy

##### Data

We downloaded model accuracy metadata and labels from CASP 8–13, and obtained the corresponding structures from the full pre-processed Protein Structure Ranking benchmarking dataset provided by the ATOM3D resource.^30^ The standard splitting methodology for accuracy estimation uses a time-based split to mirror the structure of the competition, in which the most recent targets are withheld for testing. We adopt a similar strategy, but since CASP 13 has released only two NMR structures, we augmented the test set with all NMR structures in CASP 11 and 12. We further ensured that no targets in the test set shared any chain-level CATH topology class with any structure in the train or validation sets. This resulted in a final test set of 59 targets (40 X-ray, 8 NMR, 11 cryo-EM). All non–X-ray structures were then pruned from the remaining training set targets to create an X-ray only training set of 331 targets. To prevent bias due to train set size, the X-ray structures in the training set were randomly subsampled to the same size, resulting in 287 X-ray, 42 NMR, and 4 cryo-EM structures. Five percent of each training set (22 targets, stratified by method for the mixed dataset) was removed at random to create a validation set. For training and validation sets, we randomly sample 140 decoys per target; for the test set, we include all decoys.

##### Training details

We separately trained a GVP model on each of the two training datasets. Each model used a batch size of 16 and otherwise default parameters (learning rate of 1e-4, 3 GVP-GNN layers, hidden dimension of 64), and was trained for 20 epochs using the Adam optimizer^31^ with default parameters. The final weights for each model were selected using the epoch which achieved maximum global Spearman correlation on the validation set.

#### Protein Sequence Design

##### Data

We obtained data from the 40% non-redundant dataset curated by Ingraham et al..^22^ This dataset consists of a total of 19,362 protein chains which are partitioned into train, validation, and test sets by CATH topology code. We evaluate on this test set, which contains 333 X-ray structures, 767 NMR structures, and 55 cryo-EM structures. We prune the training and validation sets as described for EMA, resulting in a training set of 13,687 X-ray structures.

##### Training details

As above, we separately trained a GVP model on each training dataset with default parameters. We trained for 150 epochs using the Adam optimizer with default parameters. The final weights for each model were selected using the epoch which achieved minimum cross-entropy loss averaged over residue predictions on the validation set.

#### Catalytic Residue Prediction

##### Data

We obtained data for functional residue prediction from the Catalytic Site Atlas (CSA).^32^ We extracted 24,836 X-ray structures and 330 non–X-ray structures. In order to prevent data leakage, we split the data into train and test splits based on sequence identity between protein chains as calculated by BLAST.^33^ 20,180 X-ray–derived chains had less than 30% sequence identity with the non-X-ray–derived chains as well as an EC class label (EC number) available. We did an 80/20 random split to create a training set (16,163 chains) and a validation set (4,017 chains). The remaining X-ray structures were set aside as a preliminary test set. The non-X-ray chains were clustered into two partitions of approximately equal size using CD-HIT.^34^ Each protein chain in the first partition had less than 30% sequence identity with each of the protein chains in the second partition, and vice-versa. One of these partitions was randomly split into a training set (160 chains) and a validation set (42 chains). The other partition (231 chains) was designated as a test set. All X-ray structures in the preliminary test set with greater than 30% sequence identity to any structure in the non–X-ray training or validation sets (1,704 chains) were removed, and the remaining X-ray structures (4,142 chains) were added to the final test set.

##### Training details

We separately trained DeepFRI on each of the two training datasets. Each model used default parameters (batch size of 64; 3 graph convolutional layers with dimensions of 128, 128, 256; dropout probability of 30%; learning rate of 2e-4; and L2 regularization with weight 1e-4), and was trained for 50 epochs using the Adam optimizer with default parameters. The final weights for each model were selected using the epoch which achieved minimum categorical cross-entropy across EC numbers on the validation set. In order to use the models for functional residue prediction, we calculated the gradient-weighted class activation maps (grad-CAMs) for each chain and predicted EC number, and used the functional importance of each residue as its predicted probability of functional activity.

## 3. Results

### 3.1. Performance on NMR and cryo-EM structures is consistently lower than performance on X-ray structures, independent of training set

First, we compared the performance of each trained model on the X-ray, NMR, and cryo-EM test sets. On all three tasks, the performance on X-ray test data was better than that of NMR and cryo-EM test data, regardless of training dataset. For EMA, both global and per-target Spearman correlations are highest for X-ray data, followed by NMR data and finally cryo-EM (Figure 2a). For protein design, the models trained on both training sets also showed significantly better perplexity on the X-ray test data than on NMR and cryo-EM data, even with a small number of cryo-EM structures (*p* < 10^− 6^; Figure 3a; Table S2). Finally, this trend is also replicated for the catalytic residue identification task. It is clear from the precision-recall curves that the model’s ability to identify functional residues is much higher for X-ray structures than for NMR structures across all recall thresholds (Figure 3b). There was not enough cryo-EM data to evaluate PR curves for this task.

**Fig. 2.**
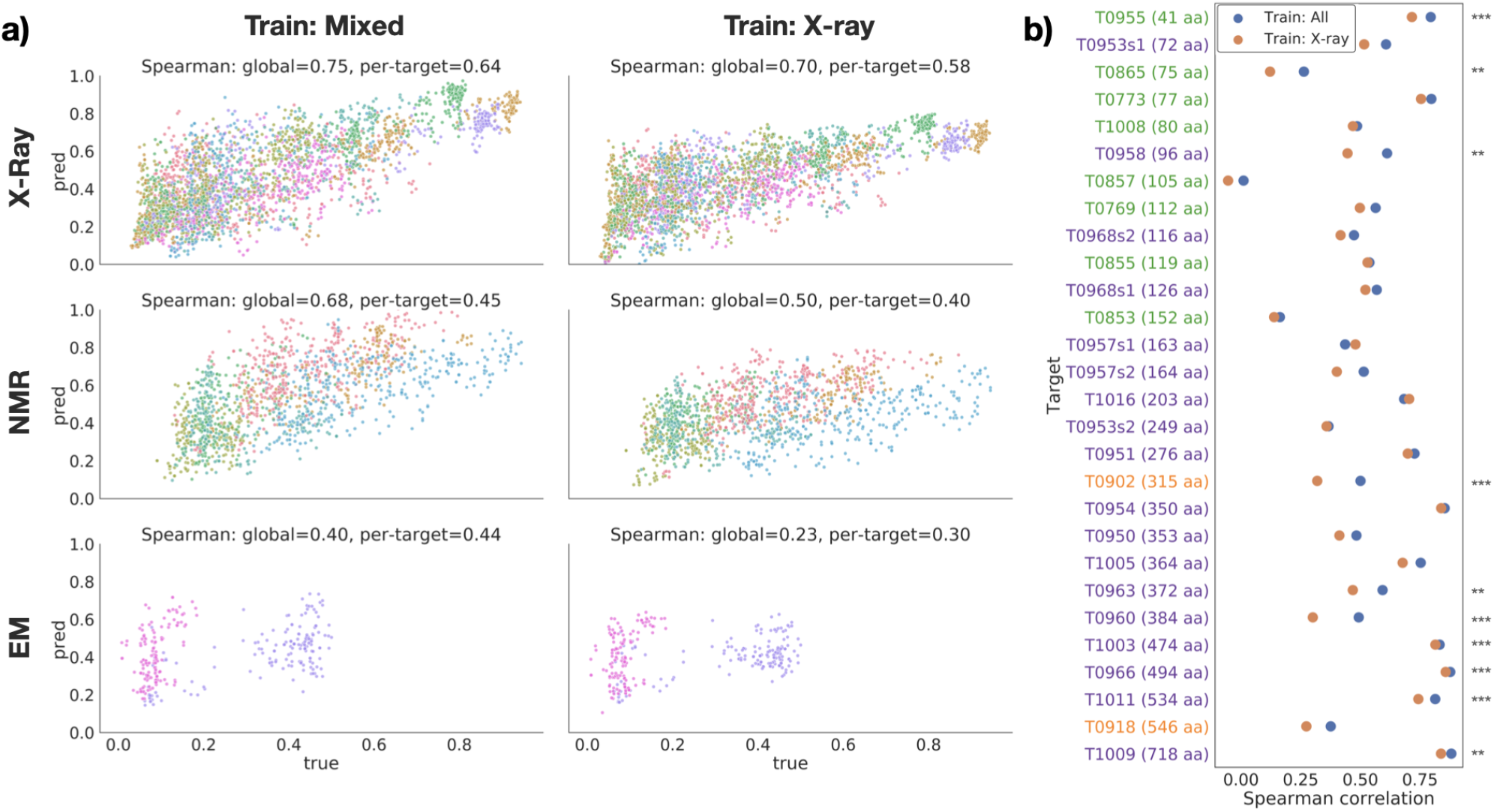
Performance on EMA task. (a) True vs. predicted GDT-TS for each target in the test set, divided by experimental method (rows), and training dataset (columns). Points are colored by target protein. (b) Per-target Spearman correlation for models trained on all structures (blue) and X-ray structures only (orange). Targets are sorted by size and colored by experimental method (blue=X-ray; green=NMR; orange=cryo-EM). Stars indicate significance for selected pairs: *** : *p* < 0.01, ** : *p* < 0.05, * : *p* < 0.1 (see Table S1).

**Fig. 3.**
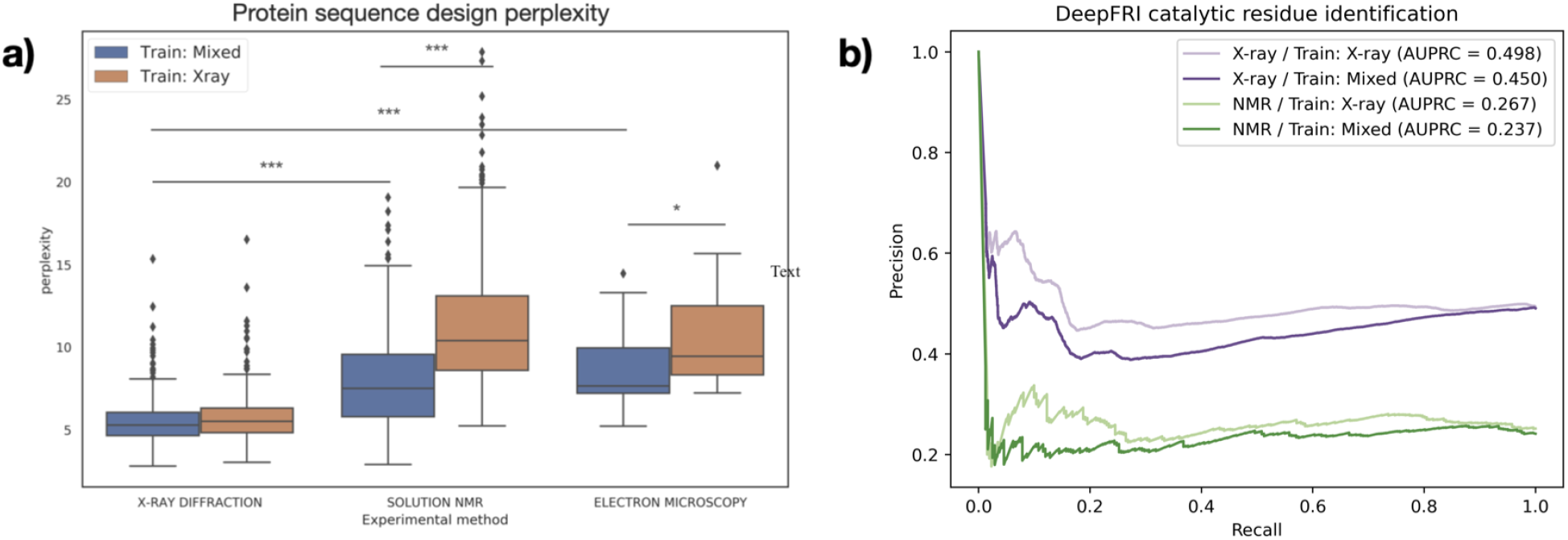
(a) Performance of GVP on protein design task test set, split by experimental method and training dataset. Lower perplexity indicates better performance. Stars indicate significance for selected pairs: *** : *p* < 0.01, ** : *p* < 0.05, * : *p* < 0.1 (see Table S2). (b) Precision-recall curves of DeepFRI on catalytic residue prediction, with X-ray test set in purple and NMR test set in green.

To ensure that the differences we observe are not due solely to differences in protein family composition between the NMR and X-ray subsets, we tested the protein design model on a set of 21 proteins with paired high-quality X-ray and NMR structures from Mei *et al* .^10^ On all of these proteins, the perplexity on the NMR structure is higher than the perplexity on the X-ray structure (mean difference of 3.71 *±* 1.63; Figure S2). Together, these data support the conclusion that models trained on PDB data are biased towards X-ray structures and the performance on NMR structures suffers as a result.

### 3.2. Inclusion of NMR data in the training set improves performance on held-out NMR data and does not degrade performance on X-ray data

Next we compared the performance of models trained on different datasets and evaluated on the same test set. In general, models trained on datasets containing all types of structure perform better on NMR test data without a loss in performance on X-ray test data. In the case of EMA, both the global and per-target Spearman correlation were higher for the GVP model trained on mixed data for all subsets of the test set (Figure 2a). The Spearman correlation for GDT-TS scores for each individual target in the test set underscores the consistency of this trend; for all but two of the 28 targets, the model trained on mixed data had a higher correlation than the model trained on only X-ray data (Figure 2b). The remaining two targets, T0957s1 and T1016, were both solved by X-ray crystallography and did not have a significant decrease in performance (*p* = 15.4739 and *p* = 20.6921; Table S1). Moreover, several of the other X-ray test structures displayed a significant increase in correlation with the inclusion of non–X-ray training data (Table S1). Figure 2b also demonstrates the differing distributions of proteins solved by each experimental method. After sorting proteins by size (i.e. number of amino acids), it is clear that NMR is used to solve much smaller structures, while cryo-EM and X-ray are used for medium to large proteins.

The GVP models for protein design also displayed this trend. While the two differently-trained models had near-identical performance on the X-ray test sets, there is a statistically significant improvement in the perplexity on the NMR test set when NMR structures are present in the training data (*p* < 10^− 90^; Figure 3a; Table S2).

One exception to this trend is the catalytic residue prediction task. While the precision-recall curves for the two models are very close to one another, the model trained only on X-ray structures slightly outperforms the model trained on a mixed dataset (Figure 3b). Because the difference in performance is not very large, this is a preliminary conclusion, though we hypothesize that this task is more suited to X-ray structures due to the high resolution required to make precise residue-level predictions.

### 3.3. Known biochemical and biophysical effects are replicated in trained models

When catalytic residue prediction models were used to predict EC number, the general trend of mixed training data improving performance held, but that of X-ray test data having higher performance than NMR test data did not. While most differences in distributions of term-centric AUPRCs were not statistically significant (Table S4), we do observe a clear upward shift in performance on NMR test data from that on X-ray test data (Figure 4a). Sorting the EC numbers into their coarser-grained enzyme classes revealed different performances across classes (Figure 4b). For example, while oxidoreductases and lyases have higher AUPRCs for X-ray test data than for NMR test data, hydrolases and translocases demonstrate the opposite. Unlike X-ray crystallography, NMR can capture the conformations in solution or associated with membranes. This may be important for broad enzyme classification. In particular, translocases are defined by their role in moving molecules across membranes, and so could be difficult to identify in a crystallized form. On the other hand, oxidoreductases commonly include catalytic metal ions, which have been historically challenging to resolve in NMR models.^35^ Because of a lack of statistical significance (Table S5), these conclusions require confirmation. Nevertheless, this demonstrates that comparing global performance may not be sufficient to evaluate machine learning models and selecting training data for specific biochemical applications.

**Fig. 4.**
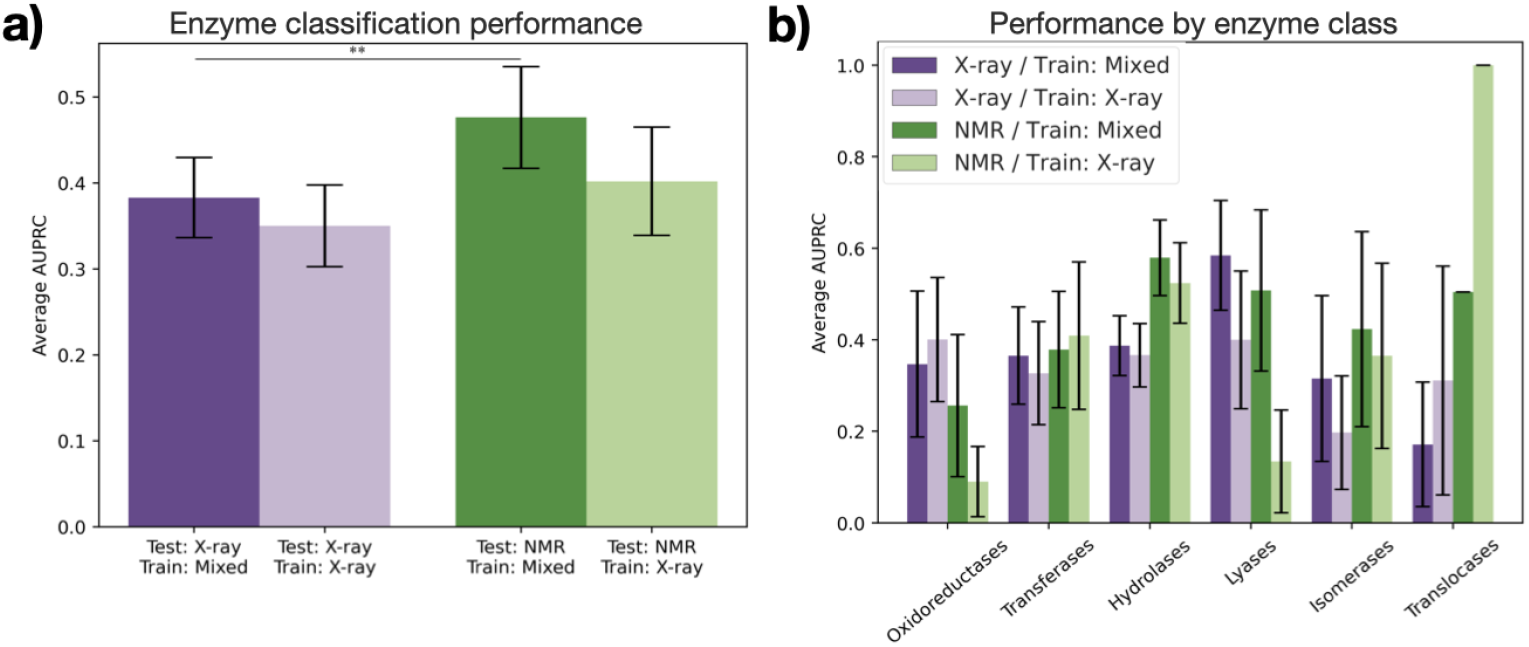
Performance of DeepFRI on enzyme classification. (a) Average AUPRC over all enzyme classes with each trained model for X-ray (purple) and NMR (green) structures in the test set. Stars indicate significance for selected pairs: ** : *p* < 0.05 (see Table S4). (b) Average AUPRC within each class of enzymes. No differences in distribution are statistically significant (see Table S5).

We further analyzed the performance of our protein design design model on specific subsets of the rest data. We first grouped the structures in the test set by secondary structure characteristics, as defined by CATH class.^36^ For proteins in the test set with mainly alpha, mainly beta, or an alpha/beta class, recovery was higher on the X-ray test structures (*p* < 0.001 except for model trained on mixed data and evaluated on mainly beta structures; Figure 5a; Table S3). These three classes also had near-identical performance on X-ray test data between the two models, but showed significant improvement for NMR test data when NMR structures were added to the training set (*p* < 10^− 8^; Table S3). Notably, many of the NMR structures in the test set with few secondary structures had higher recovery than NMR structures in other CATH classes when predicted with the model trained on mixed data, in many cases even higher than that of the X-ray structures in the test set. This is consistent with the fact that unstructured regions are often modelled better with NMR than X-ray crystallography.^37^

**Fig. 5.**
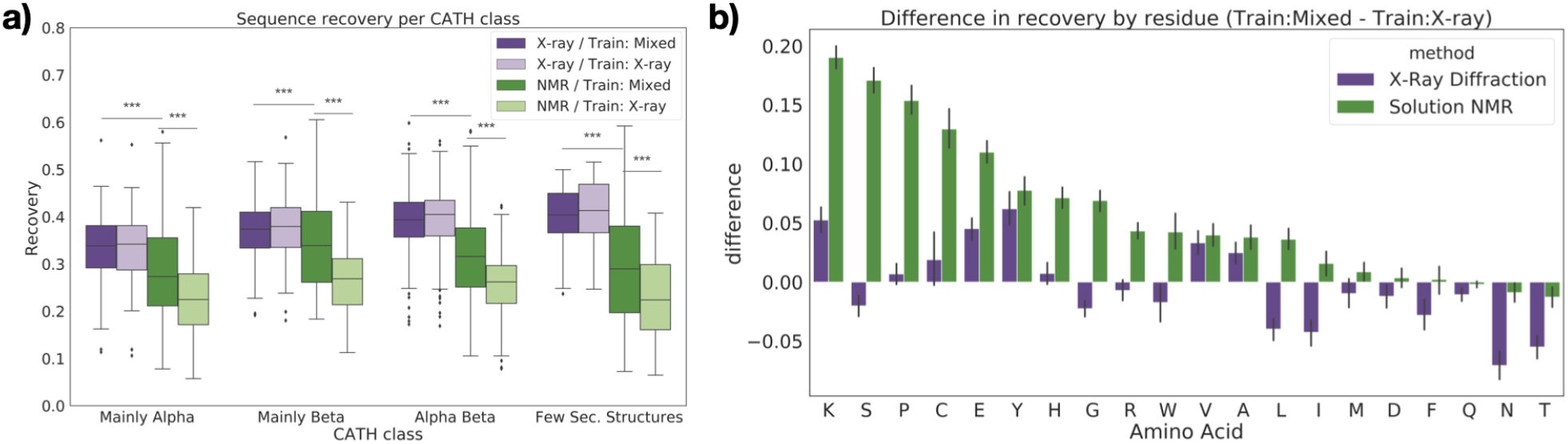
Sequence recovery for protein design as a function of protein features. (a) Sequence recovery across all residues, separated by CATH class. Stars indicate significance for selected pairs: *** : *p* < 0.01 (see Table S3). (b) Difference in recovery of each residue type when non–X-ray structures are included in the training data, separated by experimental method.

We also analyzed the performance of the protein design model in terms of its ability to predict specific amino acids. Recovery of most amino acids improved on NMR test data when NMR structures were included in the training set (Figure 5b). The greatest improvement is observed for lysine, serine, proline, cysteine, and glutamic acid. Lysine has the longest side chain of the natural amino acids, and is among the most solvent-exposed (over 90%).^38^ Lysine thus has very high conformational entropy which discourages crystallization, resulting in conformations which are captured differently in solution.^39^ Serine and glutamic acid are also polar residues with high solvent accessibility and a tendency to prefer loops and unstructured regions.^40^ Cysteine is unique in its ability to form disulfide bridges, a biochemical feature which is more difficult to resolve with NMR than X-ray crystallography.^41,42^ Core packing, which is stabilized by disulfide bridges, is also known to be significantly different between X-ray and NMR structures.^10^ Proline is known to disrupt helices and sheets, corroborating the evidence from Figure 5a that recovery improves for NMR structures with few secondary structures.

For X-ray structures, the recovery of most amino acids either improved or stayed roughly constant when trained on the mixed dataset. A confusion matrix of the errors made on these structures shows that in cases where performance does decrease, the model tends to substitute chemically similar amino acids (Figure S3, Table S6).

### 3.4. Downsampling X-ray structures during training negatively affects performance on all types of data

One logical strategy for improving the performance on NMR data would be to try to balance the training dataset by downsampling the X-ray structures. However, it is also well-known in machine learning that having a larger training dataset leads to better performance, even when that dataset may be slightly noisy. We sought to determine the nature of this tradeoff between data quality and data quantity by training our protein design model on datasets with varying degrees of downsampling (i.e. increasing class balance). The results clearly show that downsampling degrades performance on both X-ray and NMR data (Figure 6). This suggests that the benefit provided by including more data outweighs the loss in performance due to overfitting to X-ray structures.

**Fig. 6.**
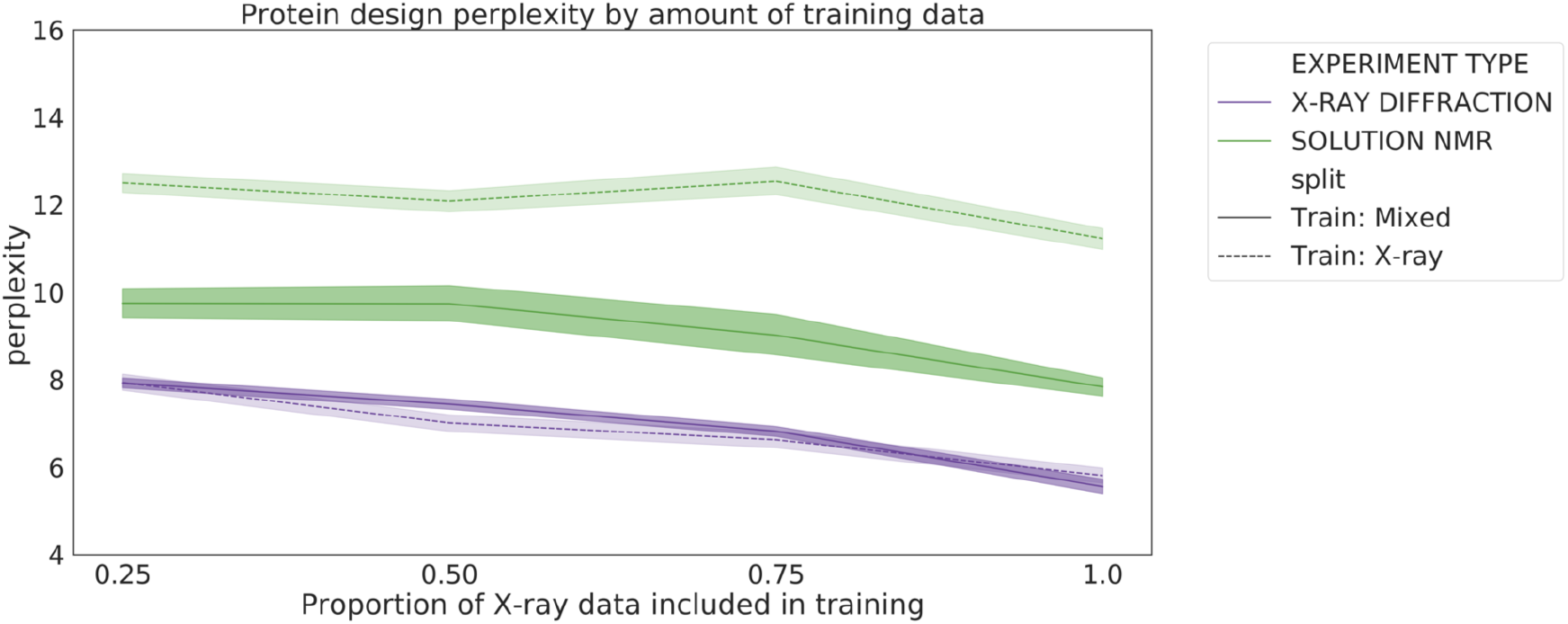
Perplexity of the protein design models as a function of training set size. Models were trained on mixed (solid line) or X-ray only (dashed line) datasets with 25%, 50%, 75%, and 100% of the X-ray training data.

## 4. Conclusion

We analyzed the impact of protein structure determination method on three machine learning tasks. This study represents, to our knowledge, the first attempt to systematically assess the effect of structure determination method on protein-based machine learning methods. Our results suggest that machine learning models perform better on X-ray crystallography data, and that performance on other types of structures are boosted by the inclusion of non–X-ray data in the training set. We show that the benefit of including more training data outweighs the penalty for overfitting or introducing noisy data for all types of test structures. However, the detailed task of interest and its sensitivity to biochemical and biophysical variables may impact the choice of data types, and global model performance may not be sufficient to measure these specific effects. Our results are preliminary, and further analysis is needed to corroborate and expand upon these findings. Our work used limited cryo-EM data, and that which was available largely had relatively low resolution. However, as the technology improves we expect to see these structures achieving similar resolution to X-ray structures. We evaluated primarily graph-based models, and so the results need to be replicated with models such as 3D convolutional networks. Finally, the recently released database of AlphaFold^2^-predicted structures provides the opportunity to train models on larger datasets than ever before. More studies are needed to assess how the inclusion of computationally-predicted structures affects the behavior of algorithms, especially for the EMA task where deep learning has notably changed the properties of decoys.^43^ Despite the need for deeper investigation, we demonstrate that in many contexts the composition of experimental methods in the training set does impact performance and should be factored into the design of machine learning methods. This work paves the way for a deeper understanding of these previously understudied factors, which will in turn enable a more principled approach to machine learning in structural biology.

## 5. Acknowledgments

We thank Dr. Joseph Puglisi and Alex Chu for discussion of experimental techniques; Dr. Xiaoyang Jing for guidance on model re-training; and Margaret Guo and Daniel Sosa for advice on experimental design and statistical interpretation. A.D. is supported by LM012409, K.A.C. is supported by LM703337, and R.B.A. is supported by NIH GM102365 and Chan Zuckerberg Biohub. Most of the computing for this project was performed on the Sherlock cluster; we would like to thank Stanford University and the Stanford Research Computing Center for providing computational resources and support.

## SUPPLEMENTARY MATERIALS

### Statistical Significance Testing

#### Estimation of Model Accuracy

Table S1 shows Bonferroni-corrected p-values for difference in GDT-TS score correlation for models trained on Mixed and X-ray only training sets. P-values are computed by applying Fisher’s transformation (see below) to convert the raw Pearson correlations to z-scores. We use Pearson for hypothesis testing because this formula is not robust for Spearman correlation. Bold indicates significance at *α* = 0.05.

Fisher’s transformation of a Pearson correlation *ρ* is computed by

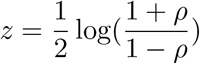

where *z* is approximately normally distributed with constant standard deviation which depends on the number of datapoints *N* :

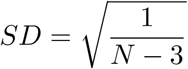

We can use the calculated *z* and *SD* to compute a two-sided p-value using a standard *z*-score hypothesis test.

#### Protein Sequence Design

Table S2 shows pairwise Bonferroni-corrected p-values for the protein sequence design task. Significance is assessed by Mann-Whitney U-test, a non-parametric test of whether two samples are drawn from the same distribution. In other words, we are testing whether there is a statistically significant shift in the distribution of protein design perplexities (1) between X-ray, NMR, and EM structures for each training set and (2) between each training set for each structure determination method. Bold indicates significance at *α* = 0.05.

#### Catalytic Residue Prediction

Table S4 shows Bonferroni-corrected p-values for the enzyme class prediction task. As above, significance is assessed by Mann-Whitney U-test. We test whether there is a statistically significant shift in the distribution of AUPRCs between the model trained on only X-ray data and the model trained on mixed data for both X-ray (row 1) and NMR (row 2) test structures. We also test whether there is a statistically significant shift in the distribution of AUPRCs between the X-ray and NMR test structures for both the model trained on only X-ray data (row 3) and the model trained on mixed data (row 4). Bold indicates significance at *α* = 0.05 after correcting for multiple hypothesis testing (*n* = 4).

Table S5 shows Bonferroni-corrected p-values for the enzyme class prediction task when the test structures are partitioned by enzyme class. As above, significance is assessed by Mann-Whitney U-test. No significant differences were found at *α* = 0.05 after correcting for multiple hypothesis testing (*n* = 24).

**Table S1.**

| Target | p-value |
| --- | --- |
| T0769 | 6.0516 |
| T0773 | 2.4567 |
| T0853 | 23.268 |
| T0855 | 19.9525 |
| T0857 | 3.1693 |
| T0865 | **0.041** |
| T0902 | 0.1559 |
| T0918 | 3.6242 |
| T0950 | 19.3462 |
| T0951 | 0.6405 |
| T0953s1 | 1.8857 |
| T0953s2 | 3.5739 |
| T0954 | 7.5116 |
| T0955 | **0.0041** |
| T0957s1 | 15.4739 |
| T0957s2 | 0.914 |
| T0958 | **0.0424** |
| T0960 | **0.0006** |
| T0963 | 0.1484 |
| T0966 | **0.0000** |
| T0968s1 | 12.3549 |
| T0968s2 | 11.3608 |
| T1003 | **0.0043** |
| T1005 | 0.6508 |
| T1008 | 11.1116 |
| T1009 | **0.0348** |
| T1011 | **0.0009** |
| T1016 | 20.6921 |

**Table S2.**

|  |  | X-ray | Train: Mixed<br>NMR | EM | X-ray | Train: X-ray<br>NMR | EM |
| --- | --- | --- | --- | --- | --- | --- | --- |
| Train: Mixed | X-ray |  | **$1.42 \times 10^{-30}$** | **$1.83 \times 10^{-5}$** |  |  |  |
|  | NMR |  |  | 1.6080 |  |  |  |
|  | EM |  |  |  |  |  |  |
| Train: X-ray | X-ray | 3.8199 |  |  |  | **$1.20 \times 10^{-123}$** | **$2.92 \times 10^{-9}$** |
|  | NMR |  | **$2.19 \times 10^{-91}$** |  |  |  | 1.7898 |
|  | EM |  |  | 0.0832 |  |  |  |

**Table S3.**

| CATH Class | Train set 1 | Test set 1 | Train set 2 | Test set 2 | p-value |
| --- | --- | --- | --- | --- | --- |
| Mainly Alpha | X-ray | X-ray | Mixed | X-ray | 10.4517 |
|  | X-ray | NMR | Mixed | NMR | **$1.11 \times 10^{-9}$** |
|  | X-ray | X-ray | X-ray | NMR | **$1.92 \times 10^{-17}$** |
|  | Mixed | X-ray | Mixed | NMR | **$1.71 \times 10^{-3}$** |
| Mainly Beta | X-ray | X-ray | Mixed | X-ray | 13.1935 |
|  | X-ray | NMR | Mixed | NMR | **$1.77 \times 10^{-13}$** |
|  | X-ray | X-ray | X-ray | NMR | **$2.24 \times 10^{-22}$** |
|  | Mixed | X-ray | Mixed | NMR | 1.9779 |
| Alpha Beta | X-ray | X-ray | Mixed | X-ray | 9.7910 |
|  | X-ray | NMR | Mixed | NMR | **$8.00 \times 10^{-20}$** |
|  | X-ray | X-ray | X-ray | NMR | **$4.36 \times 10^{-82}$** |
|  | Mixed | X-ray | Mixed | NMR | **$9.03 \times 10^{-18}$** |
| Few Secondary Structures | X-ray | X-ray | Mixed | X-ray | 16.4823 |
|  | X-ray | NMR | Mixed | NMR | **$3.90 \times 10^{-3}$** |
|  | X-ray | X-ray | X-ray | NMR | **$1.77 \times 10^{-18}$** |
|  | Mixed | X-ray | Mixed | NMR | **$2.85 \times 10^{-4}$** |

**Table S4.**

| Train set 1 | Test set 1 | Train set 2 | Test set 2 | p-value |
| --- | --- | --- | --- | --- |
| X-ray | X-ray | Mixed | X-ray | 1.2588 |
| X-ray | NMR | Mixed | NMR | 0.6264 |
| X-ray | X-ray | X-ray | NMR | 0.1880 |
| Mixed | X-ray | Mixed | NMR | **0.0404** |

**Table S5.**

| Enzyme Class | Train set 1 | Test set 1 | Train set 2 | Test set 2 | p-value |
| --- | --- | --- | --- | --- | --- |
| Oxidoreductases | X-ray | X-ray | Mixed | X-ray | 11.4456 |
|  | X-ray | NMR | Mixed | NMR | 6.3624 |
|  | X-ray | X-ray | X-ray | NMR | 3.9312 |
|  | Mixed | X-ray | Mixed | NMR | 11.3136 |
| Transferases | X-ray | X-ray | Mixed | X-ray | 10.0272 |
|  | X-ray | NMR | Mixed | NMR | 12.0000 |
|  | X-ray | X-ray | X-ray | NMR | 2.7624 |
|  | Mixed | X-ray | Mixed | NMR | 3.2688 |
| Hydrolases | X-ray | X-ray | Mixed | X-ray | 10.5024 |
|  | X-ray | NMR | Mixed | NMR | 8.6088 |
|  | X-ray | X-ray | X-ray | NMR | 0.9696 |
|  | Mixed | X-ray | Mixed | NMR | 0.3456 |
| Lyases | X-ray | X-ray | Mixed | X-ray | 3.9840 |
|  | X-ray | NMR | Mixed | NMR | 3.3408 |
|  | X-ray | X-ray | X-ray | NMR | 7.5936 |
|  | Mixed | X-ray | Mixed | NMR | 11.0880 |
| Isomerases | X-ray | X-ray | Mixed | X-ray | 8.9688 |
|  | X-ray | NMR | Mixed | NMR | 12.0000 |
|  | X-ray | X-ray | X-ray | NMR | 1.9896 |
|  | Mixed | X-ray | Mixed | NMR | 2.892 |
| Translocases | X-ray | X-ray | Mixed | X-ray | 12.0000 |
|  | X-ray | NMR | Mixed | NMR | 12.0000 |
|  | X-ray | X-ray | X-ray | NMR | 4.4520 |
|  | Mixed | X-ray | Mixed | NMR | 4.4520 |

### Relative performance on NMR structures by CASP year in test set

The only difference between our test set and the standard CASP13 dataset is that we add 6 additional NMR structures from CASP 11-12 due to the lack of NMR structures in recent CASP editions. However, the characteristics of decoys in CASP have changed over the years, especially with the advent of deep learning–based structure prediction models, resulting in potential bias when performance structures from different years are compared directly.^43^ To demonstrate that our results are consistent despite the inclusion of prior CASP data, we separate out the per-target Spearman correlations for NMR structures in each CASP in Figure S1. It is clear from this that performance decreased when training only on X-ray data regardless of which year the test structures came from.

**Fig. S1.**
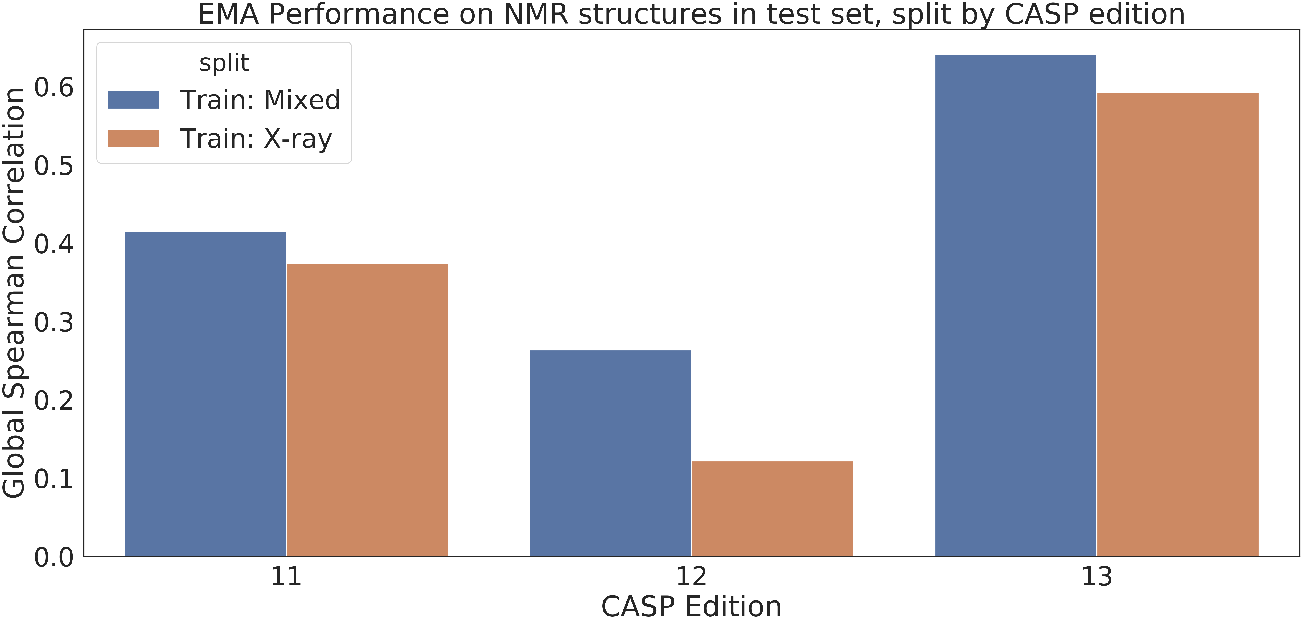
Per-target Spearman correlations for NMR structures in the test set, split out by CASP edition. Regardless of CASP edition, the correlations decrease when trained on X-ray structures only, with even greater decreases for the most recent editions.

### Per-protein protein design results for paired dataset

**Fig. S2.**
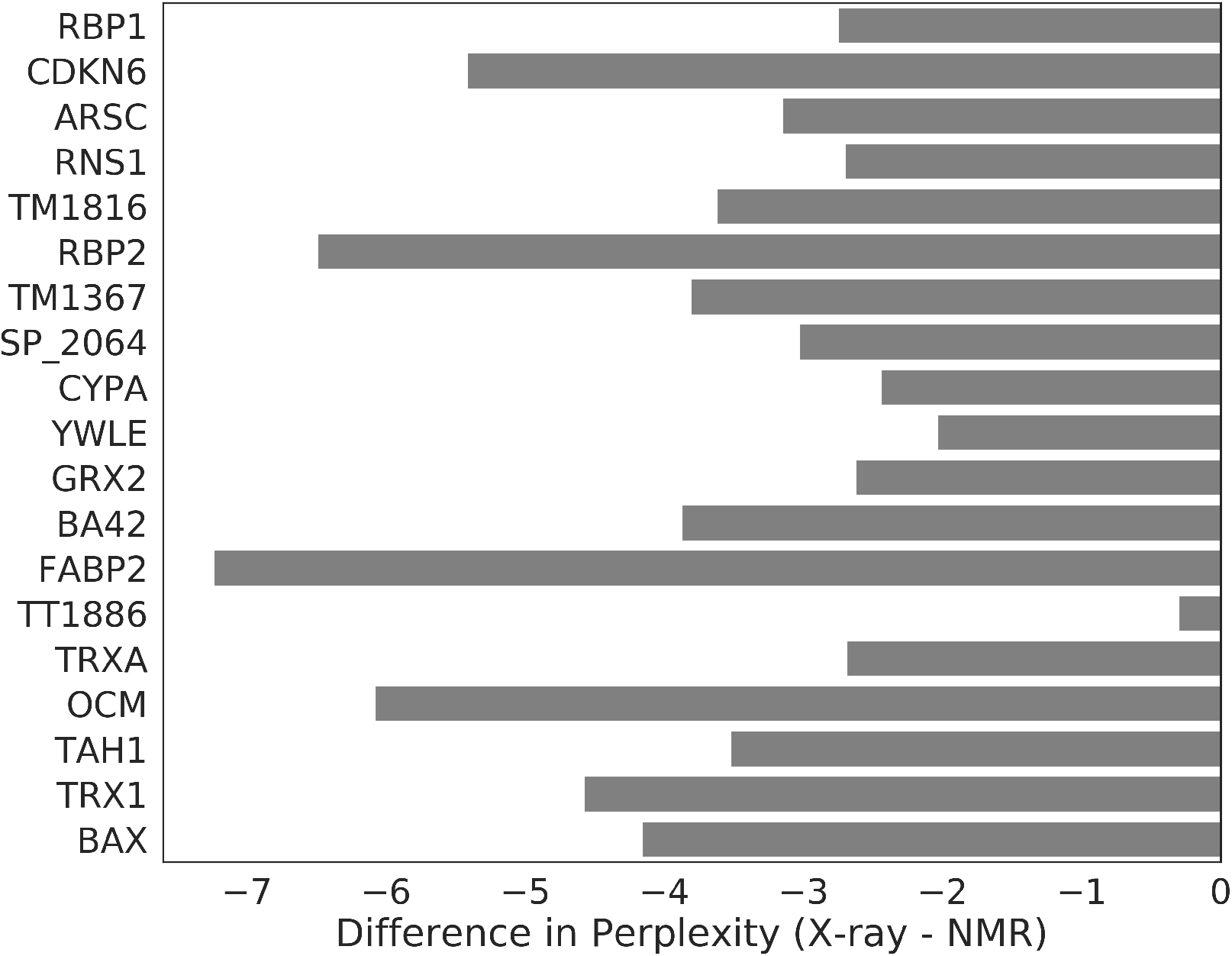
Difference in perplexity between X-ray and NMR structures for protein each protein in paired dataset from Mei et al..^10^ All proteins have lower perplexity for X-ray than for NMR structures.

### Error analysis for protein design task

Figure S3 shows the full confusion matrix for protein design on X-ray data in order to help interpret the drop in performance for certain residues. Rows are true amino acids, columns are predicted amino acids. Amino acids are ordered by biochemical properties: positively charged (H, K, R), negatively charged (D, E), small polar (S, T, N, Q), small hydrophobic (A, V, L, I, M), large hydrophobic (F, Y, W), and unique (P, G, C). Table S6 outlines most-commonly predicted residues (incorrect predictions with > 5% frequency) for the cases where performance decreased upon inclusion of NMR data in training.

**Fig. S3.**
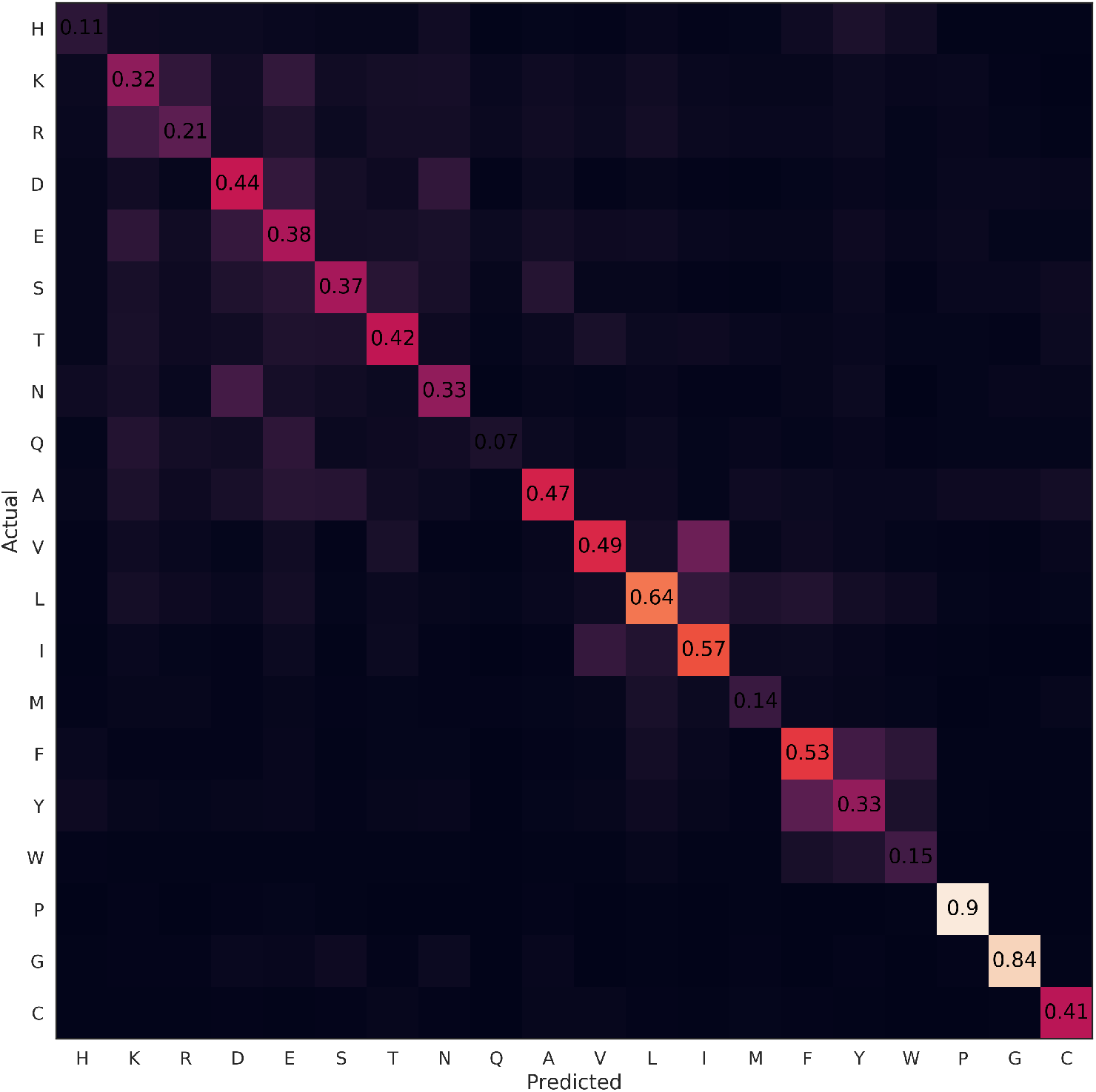
Confusion matrix for protein design task on X-ray test data. Results show that confusion is generally seen between biochemically similar residues, suggesting that the decrease in performance for certain residues may not be detrimental to model performance.

**Table S6.**

| True Residue | Predicted Residue | Frequency |
| --- | --- | --- |
| Asparagine (N) | ASP | 0.199 |
|  | GLU | 0.079 |
|  | LYS | 0.067 |
|  | SER | 0.056 |
| Threonine (T) | GLU | 0.084 |
|  | VAL | 0.076 |
|  | SER | 0.075 |
|  | LYS | 0.061 |
| Isoleucine (I) | VAL | 0.148 |
|  | LEU | 0.120 |
| Leucine (L) | ILE | 0.082 |
| Phenylalanine (F) | TYR | 0.131 |
|  | LEU | 0.104 |
| Serine (S) | ALA | 0.105 |
|  | GLU | 0.102 |
|  | THR | 0.093 |
|  | ASP | 0.067 |
|  | LYS | 0.055 |

### Dataset statistics

**Fig. S4.**
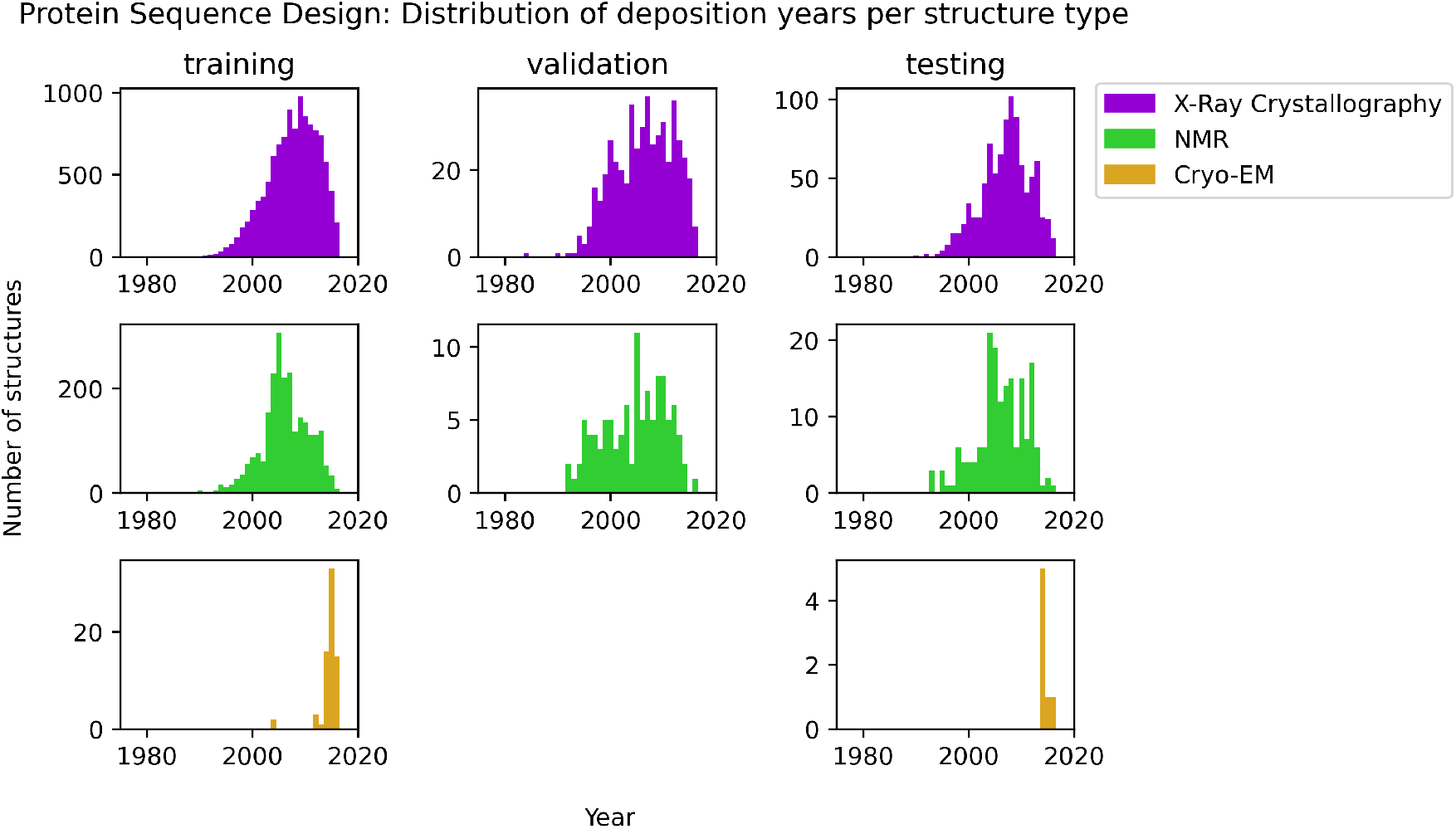
Distributions of deposition year for each structure in the training, validation, and testing sets for the protein sequence design task. Structures solved in later years tend to be higher quality than those solved in earlier years; the splitting methodology did not introduce any time bias. X-ray structures are dominant in all three splits. No cryo-EM structures were present in the validation set.

**Fig. S5.**
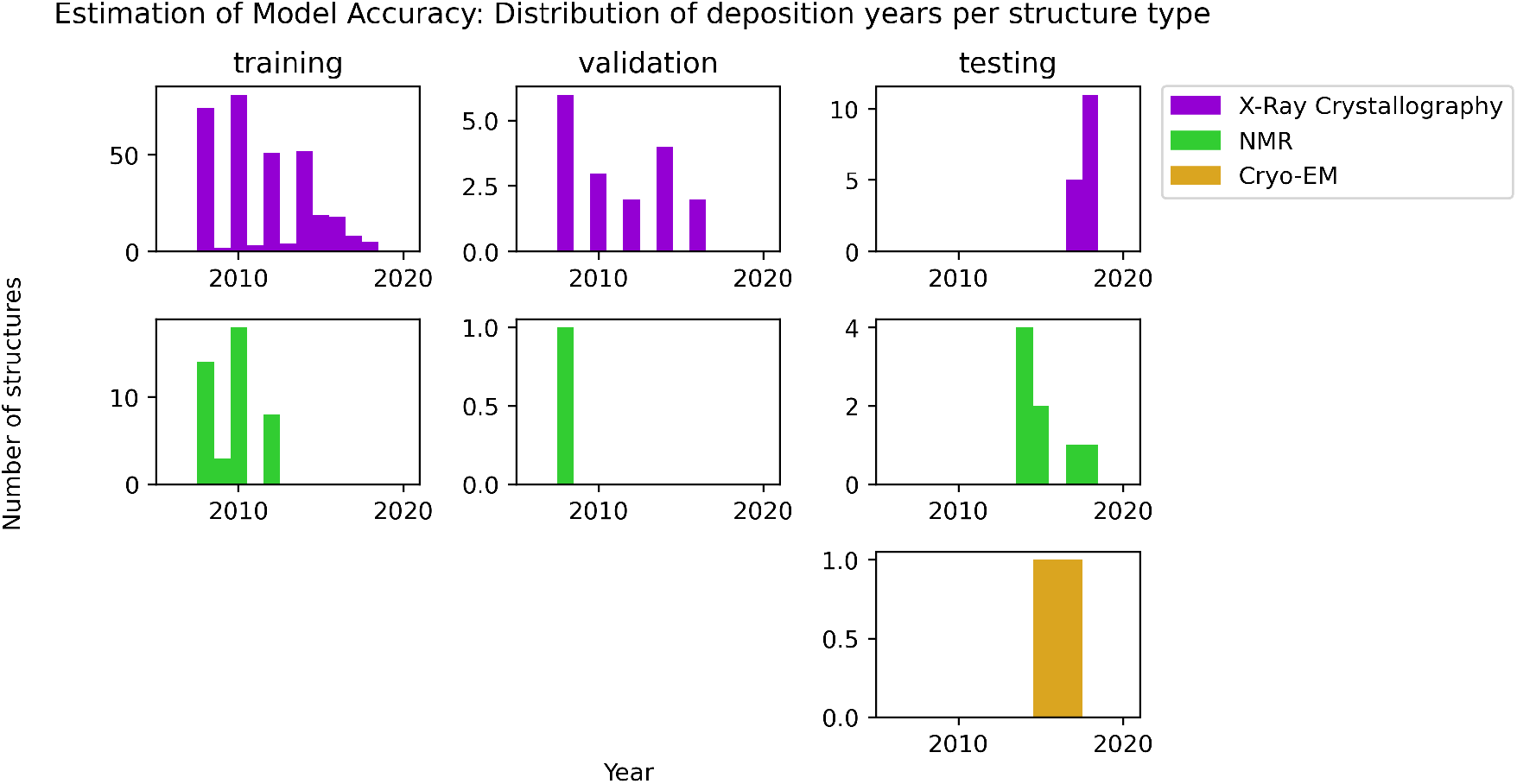
Distributions of deposition year for each structure in the training, validation, and testing sets for the evaluation of model accuracy task. As per CASP convention, a time-based split was used. X-ray structures are dominant in all three splits. No cryo-EM structures were present in the training or validation sets.

**Fig. S6.**
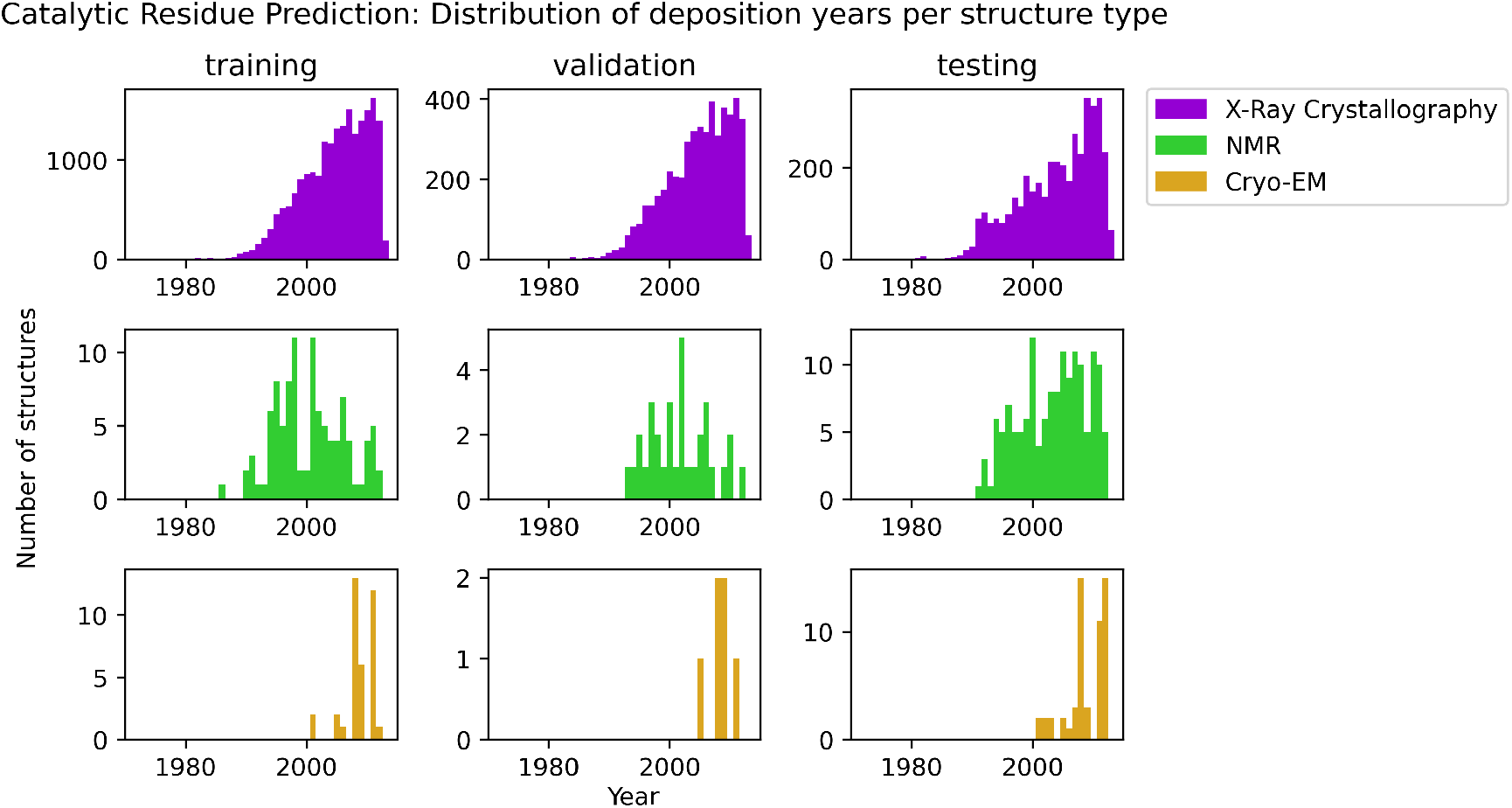
Distributions of deposition year for each structure in the training, validation, and testing sets for the catalytic residue prediction task. Structures solved in later years tend to be higher quality than those solved in earlier years; the splitting methodology did not introduce any time bias. X-ray structures are dominant in all three splits.

**Fig. S7.**
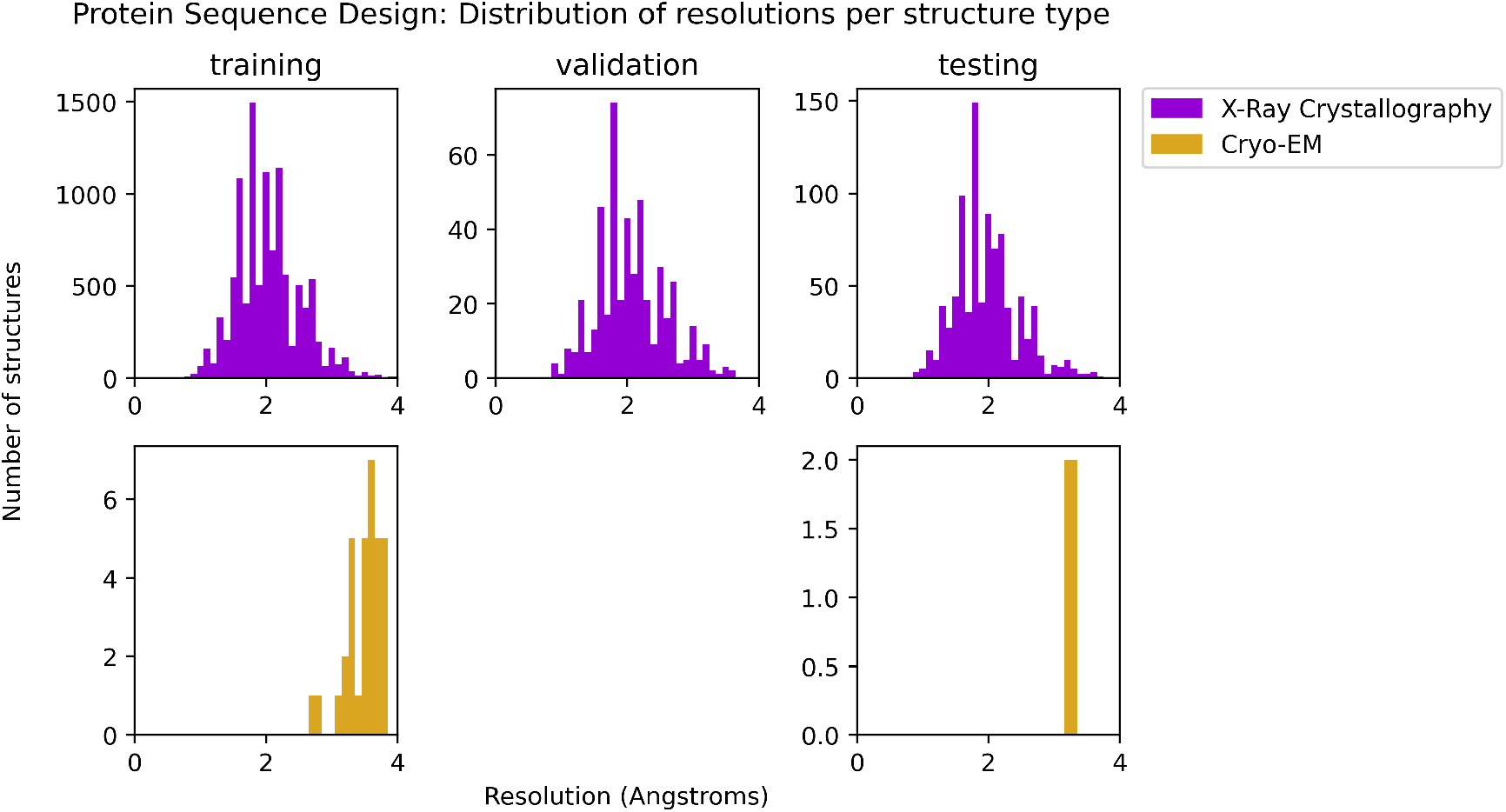
Distributions of resolution for each X-ray and cryo-EM structure in the training, validation, and testing sets for the protein sequence design task. The splitting methodology did not introduce any resolution bias. X-ray structures are dominant in all three splits. No cryo-EM structures were present in the validation set.

**Fig. S8.**
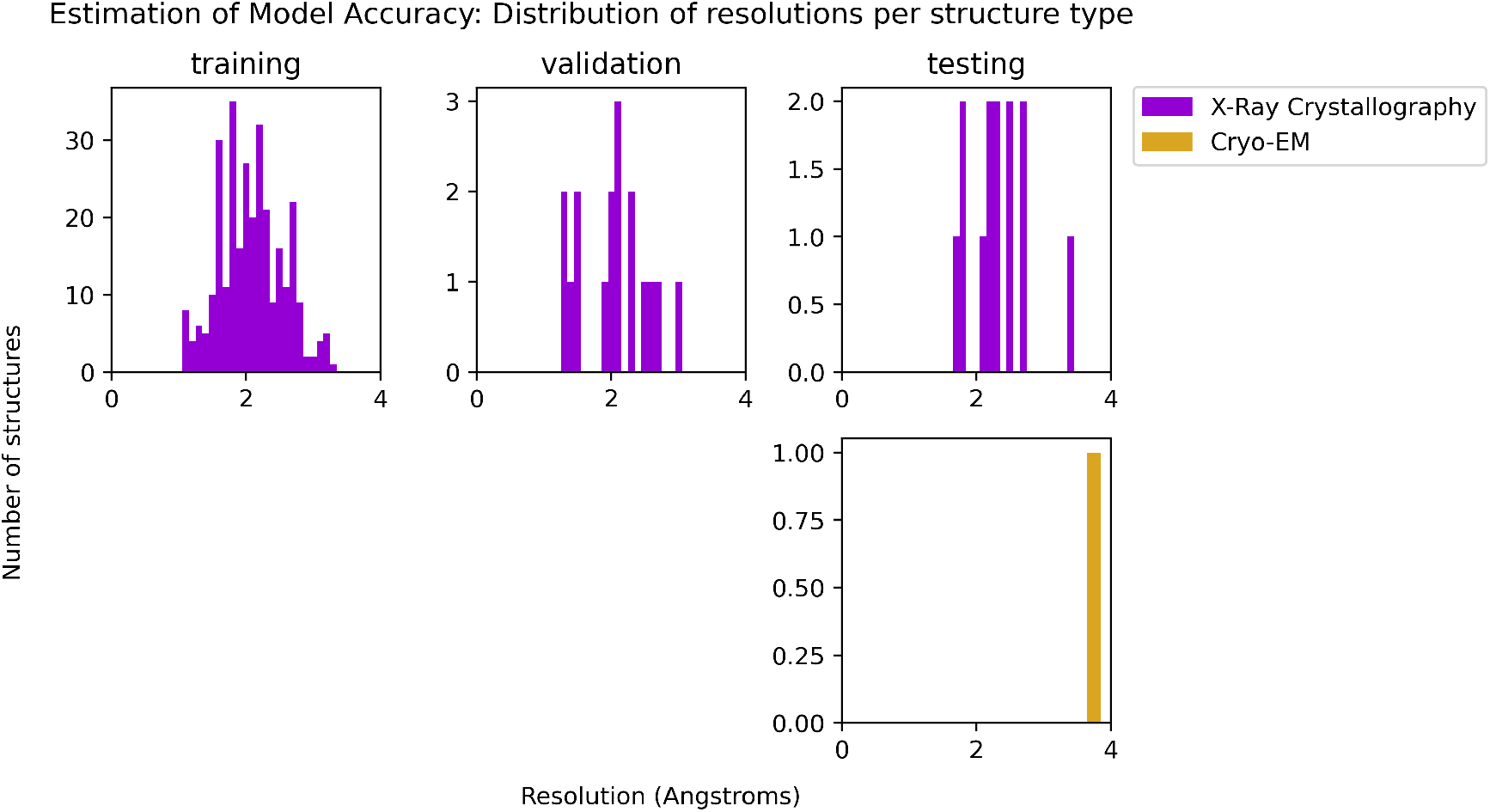
Distributions of resolution for each X-ray and cryo-EM structure in the training, validation, and testing sets for the estimation of model accuracy task. The splitting methodology did not introduce any resolution bias. X-ray structures are dominant in all three splits. No cryo-EM structures were present in the training or validation sets

**Fig. S9.**
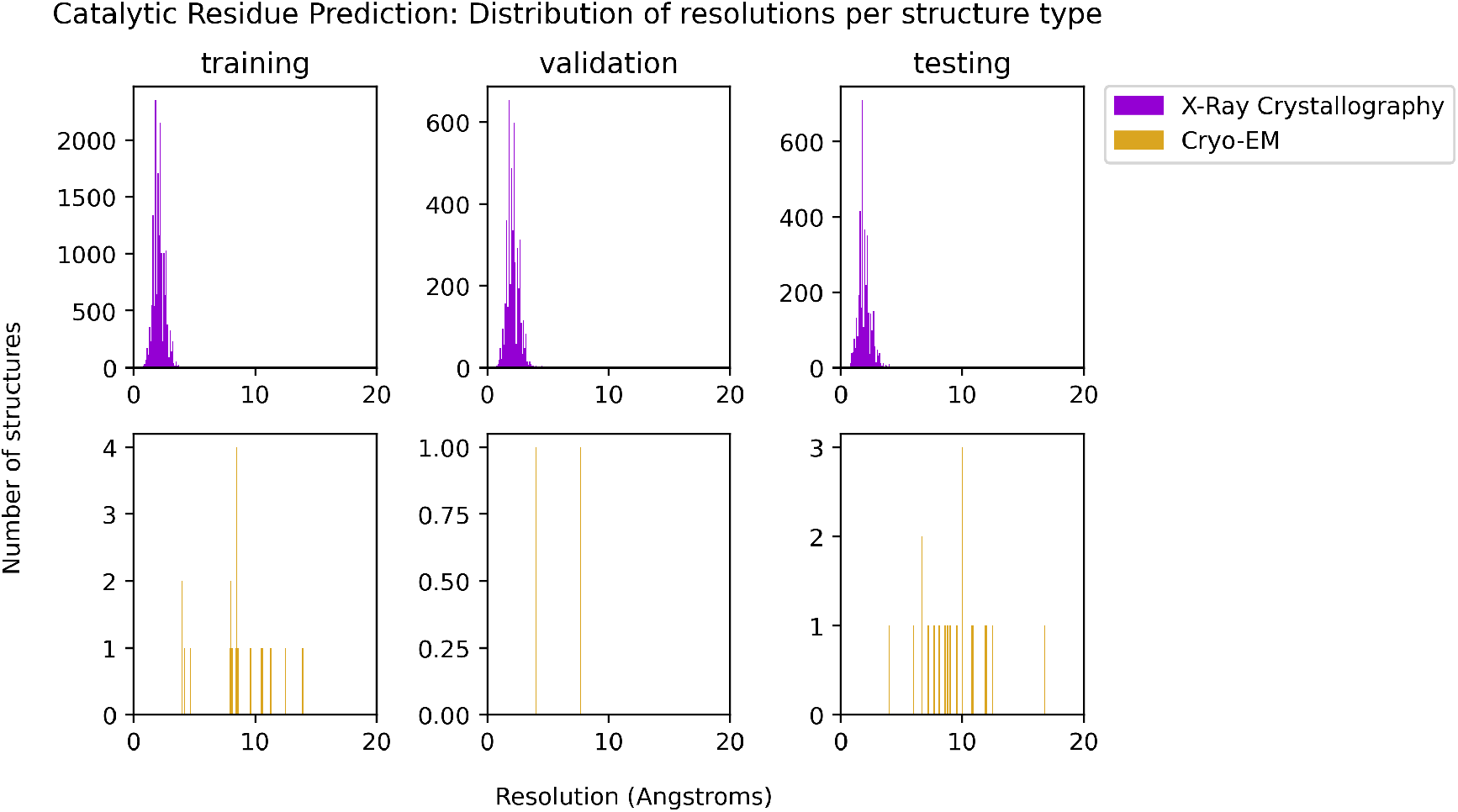
Distributions of resolution for each X-ray and cryo-EM structure in the training, validation, and testing sets for the catalytic residue prediction task. The splitting methodology did not introduce any resolution bias. X-ray structures are dominant in all three splits.

## Notes

### Competing Interest Statement

The authors have declared no competing interest.

## References

1. M. Baek, F. DiMaio et al., Science 10 (2021).

2. J. Jumper, R. Evans et al., Nature (2021).

3. J. Yang, I. Anishchenko et al., PNAS 117 (2020).

4. R. Townshend, R. Bedi et al., NeurIPS 32 (2019).

5. P. Gainza, F. Sverrisson et al., Nat. Methods 17 (2020).

6. N. Anand and P. Huang, NeurIPS (2018).

7. C. Norn, B. I. Wicky et al., PNAS 118 (2021).

8. N. Anand-Achim, R. R. Eguchi et al., bioRxiv (2021).

9. L.-W. Yang, E. Eyal et al., Structure 15 (2007).

10. Z. Mei, J. D. Treado et al., Proteins Struct. Funct. Bioinf. 88 (2020).

11. S. O. Garbuzynskiy, B. S. Melnik et al., Proteins: Struct. Funct. Genet. 60 (2005).

12. M. Andrec, D. A. Snyder et al., Proteins: Struct. Funct. Genet. 69 (2007).

13. V. Krishnan and B. Rupp, eLS (2012).

14. H.-W. Wang and J.-W. Wang, Protein Sci. 26 (2017).

15. M. A. Marques, M. D. Purdy and M. Yeager, Curr. Opin. Struct. Biol. 58 (2019).

16. J. Quiñonero-Candela, M. Sugiyama et al. (MIT Press, 2009).

17. H. Berman, K. Henrick et al., Nucleic Acids Res. 35 (2007).

18. A. Kryshtafovych, T. Schwede et al.

19. A. Zemla, Nucleic Acids Res. 31 (2003).

20. J. Wang, H. Cao et al., Sci. Rep. 8 (2018).

21. J. O’Connell, Z. Li et al., Proteins 86 (2018).

22. J. Ingraham, V. K. Garg et al., NeurIPS (2019).

23. B. Jing, S. Eismann et al., 2009.01411 (2020).

24. B. Kuhlman and D. Baker, PNAS 97 (2000).

25. N. Hulo, A. Bairoch et al., Nucleic Acids Res. 34 (2006).

26. M. Blum, H.-Y. Chang et al., Nucleic Acids Res. 49 (2021).

27. B. Jing, S. Eismann et al., 2106.03843 (2021).

28. V. Gligorijević, P. D. Renfrew et al., Nat. Commun. 12 (2021).

29. A. Bairoch, Nucleic Acids Res. 28 (2000).

30. R. J. Townshend, M. Vögele et al., 2012.04035 (2020).

31. D. P. Kingma and J. Ba, 1412.6980 (2014).

32. N. Furnham, G. L. Holliday et al., Nucleic Acids Res. 42 (2014).

33. S. F. Altschul, W. Gish et al., J. Mol. Biol. 215 (1990).

34. L. Fu, B. Niu et al., Bioinformatics 28 (2012).

35. D. K. Chakravorty, B. Wang et al., J. Biomol. NMR 56 (2013).

36. C. A. Orengo, A. D. Michie et al., Structure 5, 1093 (1997).

37. T. Rehm, R. Huber and T. A. Holak, Structure 10, 1613 (2002).

38. F. Baud and S. Karlin, PNAS 96, 12494 (1999).

39. Z. S. Derewenda and P. G. Vekilov, Acta Crystallogr. D Biol. Crystallogr. 62, 116 (2006).

40. A. K. Shaytan, K. V. Shaitan and A. R. Khokhlov, Biomacromolecules 10, 1224 (May 2009).

41. B. Schmidt and P. J. Hogg, BMC Struct. Biol. 7, 1 (2007).

42. D. A. Armstrong, Q. Kaas and K. J. Rosengren, Chem. Sci. 9, 6548 (2018).

43. J. Won, M. Baek et al., Proteins Struct. Funct. Bioinf. 87, 1351 (12 2019).

